# Joint ancestry inference reveals the landscape of archaic introgression in admixed populations

**DOI:** 10.64898/2026.08.29.748036

**Authors:** Jazeps Medina Tretmanis, Valeria Añorve-Garibay, David Peede, Mayra M. Bañuelos, María C. Ávila-Arcos, Flora Jay, Emilia Huerta-Sanchez

**Author notes:** **Corresponding authors:**, emilia. Contributed equally (María C. Ávila-Arcos, Flora Jay, Emilia Huerta-Sanchez).

## Abstract

Studying the evolutionary history of archaic segments in recently admixed individuals requires inferring both continental and archaic ancestry in admixed genomes. Here, we present *TRACTINATOR*, the first deep-learning method for simultaneous inference of continental and archaic ancestry in admixed human genomes. The model combines SNP sequences, population allele-frequency information, and S* statistics to improve both inference tasks. By learning relationships between haplotypes and population allele frequencies, *TRACTINATOR* can generalize across genomic regions and even across different genomic datasets. We train our model using both real and synthetic data, and show that augmenting with synthetic data improves accuracy for both continental and archaic ancestry inference. Finally, we apply *TRACTINATOR* to admixed Latin American populations from the 1,000 Genomes Project, revealing how archaic ancestry is distributed within chromosomal segments of African, European and Indigenous American ancestry in Latin American individuals. For candidates of adaptive introgression, we also infer whether the archaic haplotype was introduced via European or Indigenous American ancestors.

## 1 Introduction

Local Ancestry Inference (LAI) has long been a cornerstone of population genomics, providing insights into the evolutionary history of recently admixed populations by identifying the ancestral origin of genomic segments. Similarly, the inference of introgressed archaic segments in modern human genomes has illuminated how historic interbreeding events with Neanderthals and Denisovans may have conferred fitness benefits as modern humans populated the globe[1][2][3]. Although both tasks aim to assign ancestry labels to genomic segments, they differ in the temporal resolution of the ancestry they seek to infer. Most LAI methods infer ancestry introduced by recent admixture among modern human populations[4][5][6], whereas archaic introgression inference targets ancestry inherited from much older gene flow events with archaic human populations[7][8][9]. Most LAI methods rely on probabilistic models, such as hidden Markov models (HMMs), and require well-characterized reference panels for each ancestral population[10]. In contrast, methods for detecting archaic introgression can be divided into two broad classes[9]: 1) reference-based approaches that leverage sequenced archaic genomes[11][12][13] and 2) reference-free approaches, which detect introgression without relying directly on sequenced archaic hominids[14][15][8].

These methods have enabled the inference of local ancestry in recently admixed individuals, providing insights into the timing and demographic history of recent admixture[16][17], patterns of sex-biased gene flow[18], and helped develop ancestry-specific genome-wide association[19] and finemapping studies[20]. Similarly, methods for detecting archaic introgression have enabled studies to identify and quantify introgressed sequences and to uncover candidates for adaptive introgression in human populations[2][21][1]. However, recently admixed individuals harbor both recent continental ancestry and archaic introgression, requiring both sources of ancestry to be considered simultaneously. In this context, recent admixture may influence the accuracy of archaic ancestry inference, while archaic introgression may likewise affect the inference of local ancestry tracts. Consequently, methods designed to infer either source of ancestry in isolation may not perform optimally in recently admixed genomes. Addressing how recent admixture has reshaped patterns of archaic ancestry therefore requires methods that jointly infer recent continental admixture and archaic introgression in modern admixed individuals. A recent method that tackles a related problem is DAIseg[15], an HMM model that can classify genomic segments as belonging to either recently admixed populations or being of archaic origin. However, it does not label both continental ancestry and archaic introgression for the same genomic segment.

Here, we present the first deep-learning method (*TRACTINATOR*) capable of jointly inferring continental and archaic introgressed regions in admixed human genomes, broadening our ability to investigate the distribution, evolution, functional and phenotypic impact of archaic ancestry in recently admixed individuals. We find that including a mixture of real and synthetic data improves the accuracy of both the continental and archaic inference tasks. Conceptually, *TRACTINATOR* is similar to the LAI approach proposed by SegNet[22] and SALAI-Net[23], training a single deeplearning model to perform inference for genomic windows at any position along the genome. One key difference is that our method trains on not just the reference haplotypes themselves, but rather on the reference haplotypes along with the alternative allele frequencies (AAFs) at each genomic position available for the source populations. This means that the model learns to identify ancestries based on similarities between a target haplotype’s SNP sequence and the AAFs of the source populations, rather than directly on the alleles in the target haplotype’s SNP sequence. One advantage of our method is that it can be trained in one genomic region and infer local ancestry in a completely different region. Another advantage is that it can be trained on one set of individuals to perform inference using AAF data derived from a different set of reference individuals.

For the task of detecting archaic introgression, we infer archaic ancestry by calculating the distribution of *S\**[7] statistics within windows along the genome. We feed summaries of this distribution to the model along with the data features necessary for continental LAI (*i.e.*, SNP sequence and AAF information). We find that supplementing *S\** information not only allows us to perform both ancestry inference tasks at the same time, but also improves the accuracy of continental LAI. Similarly, the inclusion of LAI features improves the accuracy of the archaic introgression task. This positions our approach alongside established reference-free tools while extending their functionality to joint inference of modern and archaic ancestry. Finally, deep-learning models require extensive training data, but the limited availability of archaic genomes and the scarcity of suitable datasets for understudied populations, such as those in Latin America, remain major challenges. To address this, we show that training our model on a mixture of the *1KG* dataset and synthetic data yields high inference accuracy on datasets that were not used for training.

We apply our method to jointly infer continental and archaic ancestry in admixed individuals from the 1,000 Genomes Project (*1KG*). Using these inferences, we characterize how recent admixture has shaped the ancestry-specific distribution of archaic segments in populations from the Americas by quantifying the prevalence of Indigenous American or European genomic regions harboring archaic tracts. Furthermore, we identify candidate regions of adaptive introgression inherited by admixed individuals through either Indigenous American or European ancestors. This work not only introduces methodological innovation in how we train and apply deep learning models in population genetics, but also provides a useful new method to jointly characterize continental and introgressed ancestry in recently admixed individuals. We showcase the strength of our approach by generating new ancestry maps for admixed individuals in the *1KG* dataset.

## 2 Methods

### 2.1 Model Input

We infer the genetic ancestry of sections of a chromosome by partitioning it into windows of 1,000 SNPs each. Each genomic window *W* is encoded as a two dimensional matrix *G_W_ ∈* R^(^*^c^*^+1^*^,s^*^)^, where *c* corresponds to the number of source populations, which is the same as the number of target classes for the model, and *s* corresponds to the number of SNPs that are included in each window.

More specifically, *G_W_* = (*g_ij_*) with *i ∈* 1, 2*, …, c*+1, and *j ∈* 1, 2*, …, s*. For *i ≤ c*, *g_ij_* corresponds to the AAF in the *i*-th population for the *j*-th SNP. When *i* = *c* + 1, *g_ij_* will be 0 if the query haplotype has the reference allele in the *j*-th position and 1 if it is the alternative allele. There are two advantages by including AAFs in this input matrix. First, it allows us to encode the target haplotype and, through AAFs, a representation of what genetic variation is common and rare in the different source populations. Second, by providing AAF information as part of the input, our model can infer the ancestry of any 1,000 SNP window in a genome, irrespective of its physical position or whether it overlaps with windows used for training.

In addition to calculating the AAFs within a genomic window *W*, we also calculate its *S\** score distribution. Specifically, for each window *W*, we calculate *S\** scores and the number of alleles absent in the outgroup population (Yoruban individuals in the *1KG*) in sliding windows of 50kb[7] with a step size of 10kb (see supplementary Section S5). We note that while each genomic window *W* contains 1,000 SNPs, different windows will vary in physical length, as expressed in base pairs. This means that given two genomic windows *W*_1_ and *W*_2_, corresponding to different physical lengths, a different number of *S\** scores will be calculated. In order to ensure that all windows have the same number of features to feed into the model, we take the mean, standard deviation, and maximum of the *S\** score. Similarly, we supply the mean, standard deviation, and maximum of the number of alleles absent in the outgroup (see supplementary Section S5). These six values are then used as features for the model. These features are not included in the input matrix *G_W_*, and instead are appended to intermediate embeddings before the final classification step of the model (see Section 2.3, Figure 1).

**Figure 1:**
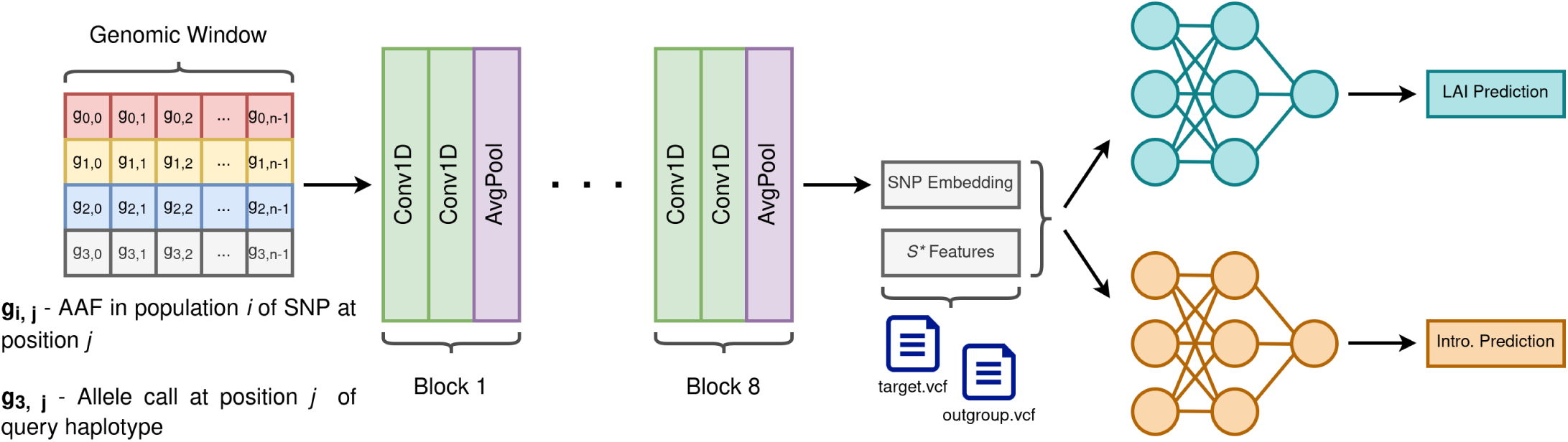
Schematic view of the model architecture. Inputs are formatted as a 2*D* matrix which includes both SNP and AAF information for a single genomic window. Embeddings of this input are produced by 8 Convolutional layers. Finally, these embeddings are supplemented with *S\** information and fed into two task-specific feedforward neural networks, resulting in one continental ancestry call and one introgression presence call.

### 2.2 Model output

The model outputs a pair of predictions for any given input matrix (see Section 2.1). One prediction is a vector of length *c*. Each entry in the vector corresponds to the probability of each possible source ancestry, and all entries sum up to 1. We only keep the ancestry (*i.e.*, the index) with the maximum probability from this vector. The other prediction is a single scalar *a,* 0 *≤ a ≤* 1, which denotes the model’s certainty of the presence of an archaic introgressed segment for the corresponding input window.

When performing the final inference on our testing and inference datasets (described in Section 2.4), we feed data into the model using overlapping windows. We use a window length of 1,000 SNPs, along with a step size of 300 SNPs. This means that each 300 SNP-region has *∼* 3 associated pairs of predictions for continental and archaic genetic ancestry. These 3 pairs of predictions are collapsed into a single pair by taking a majority vote of the continental ancestry predictions, and taking the mean of the 3 archaic ancestry predictions. We then discretize the mean archaic ancestry probabilities into a binary label, taking values larger than 0.5 as a positive label for archaic introgression. After combining overlapping inferences, the final outputs are continental and archaic labels for genomic regions of 300 SNPs, which average 14.13kbp *±* 1.96kbp in length in the *1KG* dataset. It should be noted that the model does not output a prediction for the donor population of archaic introgressed tracts (e. g., Neanderthal or Denisovan).

### 2.3 Model architecture

The neural network is structured as a Convolutional Neural Network (CNN) encoder that transforms the input matrix *G_W_*(see Section 2.1) into a vector of SNP encodings *E_W_*. The input matrices are passed through sequential blocks of 1*D* convolutional layers, with an initial input channel count of *c* + 1, with *c* being the number of reference populations considered (the extra channel being used for the target SNP sequence itself). Each 1*D* convolution is followed by batch normalization and rectified linear unit (ReLU) activation. Progressive channel expansions (8, 16, 32, 64, 128, 256, and 512 channels) are introduced across 21 convolutional layers, with average pooling applied after every two convolutional layers to reduce feature dimensionality. This feature extractor outputs *E_W_*, the fixed-length encoding for the input matrix *G_W_*. A detailed description for each layer is provided in Section S2 of the supplementary material.

This vector *E_W_* is then concatenated with the mean, standard deviation and maximum of the *S\** score and non-outgroup SNP count distributions for window *W* (Figure 1). The combined representation is passed to two task-specific fully-connected classifiers. For the continental ancestry (LAI) task, three stacked fully-connected layers (each of size 2,054 with ReLU activations and dropout rate = 0.2) precede a final classification softmax layer outputting a vector *v* of length *c*. Each of these *v_i_* values represents the probability that the reference population *i* is the continental ancestry for window *W*. For the archaic ancestry task, a similar multilayer perceptron outputs a single value that corresponds to archaic introgression probability. When training the model, we calculate the cross-entropy loss for both tasks independently; these two loss values are added together before performing the backpropagation step. We do not transform or scale the losses before adding them together, as they are close enough in scale that adding them together does not bias the model towards any particular task.

### 2.4 Datasets

Here, we describe the testing and training data sets used for our joint continental and archaic ancestry caller. We consider multiple combinations of simulated data (using *msprime* v.1.2.0[24]) and genomic data from unadmixed populations from the *1KG* panel. Specifically, we train exclusively on synthetic data or on a combination both synthetic data and *1KG* data. For testing, we benchmark our model using synthetic data alone or a combination of synthetic and real data from the *1KG* or Simons Genome Diversity Project[25] (*SGDP*) panels.

#### 2.4.1 Synthetic training and testing data

We use *msprime* to simulate synthetic genomes that we then partition into non-overlapping windows to use as training data. These genomes are simulated using three commonly used demographic models that have been used to simulate admixed individuals from the Americas[26][21] (illustrated in supplementary Section S3). In particular, we sample simulated African (AFR), European (EUR), and, depending on the demographic model, an East Asian (EAS, models *A* and *B*) or Indigenous American (MXB, model *C*) population to use for training. For the testing data, we sample an admixed population (ADMIXED) with genetic contributions from AFR, EUR, and either EAS or MXB to mimic admixture and introgression patterns in the Americas at varying degrees of realism. The specific demographic models used to simulate the training data are:

A. Admixture model in the Americas described by Browning et al. (2018)[26].
B. Admixture model in the Americas described by Browning et al. (2018)[26], modified to include a single pulse of introgression of a Neanderthal lineage into the Eurasian branch 1, 600 generations in the past.
C. Admixture model in the Americas described by Villanea and Peede et al. (2025)[21], modified to simulate both a Neanderthal and a Denisovan pulse of introgression.

The demographic model *A* does not simulate introgression, while models *B* and *C* do include introgression from Neanderthals or Denisovans. For each demographic history, we simulated 700 haploid genomes (20Mb long) that were partitioned into non-overlapping genomic widows containing 1,000 SNPs each. For each simulation, we sampled unadmixed synthetic individuals from AFR (*n* = 300), EUR (*n* = 200) and EAS (*n* = 200) populations for training. The reason we sample more AFR individuals than other ancestries, is that 100 of these synthetic AFR individuals are only used for *S\** calculations (see Section 2.1). The remaining 600 unadmixed individuals (200 individuals from each population), along with their respective *S\** score and non-outgroup SNP count distributions, are used for model training.

We always use a recombination map of chromosome 1 when simulating these synthetic individuals[27]. For model *B*, we vary the introgression proportion (values of 1%, 2%, 5%, 10%, 15%) and simulate one replicate (700 haploid, 20Mb genomes) for each proportion. This is useful as it allows the model to generalize on different SNP configurations from those available only in real training data. For model *C*, we do not vary the introgression proportion. All demographic parameters for models *A, B* and *C* and counts of simulated windows are listed in Section S3 of the supplementary material. Since genomic windows are used as training data, it is necessary to extract the ground truth continental and archaic ancestry labels for each individual at each window. We use the resulting tree sequences from *msprime* to obtain the true labels (for archaic and continental ancestry) in the simulated data. The algorithm for labeling of introgressed windows is described in Section S4 of the supplementary materials.

In total, across all demographic models, we use 10,000 non-overlapping genomic windows of synthetic training data by sampling the unadmixed populations. The entire process is repeated to generate 10,000 non-overlapping genomic windows of synthetic testing data by sampling the admixed populations (ADMIXED *n* = 200) and obtaining AAFs for the AFR, EUR and EAS populations in the simulation. We note that even though we are using identical parameters for these testing individuals, the SNP positions and allele frequencies of the testing and training windows are different due to the inherent randomness of the simulations.

#### 2.4.2 Training and testing data from the 1KG

When training and testing the model on real data, we use three unadmixed populations from the *1KG* dataset. AFR (excluding YRI), EUR, and EAS, which combined result in 1,452 unadmixed individuals. For every population, we take 90% of the available individuals to use for training, resulting in 1, 296 unadmixed training individuals. To compute *S\** scores (Section 2.1), we use YRI individuals (*n* = 108) that are excluded from the training and testing datasets. YRI is the outgroup population required to compute *S\** scores (Section 2.1), as they represent a population assumed to lack an appreciable amount of Neanderthal or Denisovan introgressed ancestry[9]. To evaluate the performance of the method, we only use windows from chromosome 1 for our training dataset, partitioning it into non-overlapping windows of 1,000 SNPs each. This results in *∼* 6, 000 genomic windows being available as training data for real individuals.

We generate two testing datasets from the *1KG* using chromosomes 1 and 22. Our inclusion of two chromosomes allows us to test two scenarios: (i) testing and training data are sourced from the same chromosome, and (ii) they are sourced from distinct chromosomes. To create these testing datasets, we use 10% of the original available unadmixed individuals, resulting in a smaller set of 156 individuals (AFR without YRI *n* = 48, EUR *n* = 45, EAS *n* = 63). Since these are unadmixed individuals, we perform a forward-in-time admixture simulation between these 156 individuals using *haptools*[28] and a human recombination map of chromosome 1 or 22[27]. We simulate 25 generations of admixture with initial proportions of AFR= ⅙, EUR= 3/6, EAS= 2/6 and generate 200 admixed genomes. When inferring the ancestry of the testing admixed individuals, we use the AAFs from the unadmixed testing individuals.

Although we have access to the continental ancestry labels for these individuals, it is impossible to have an empirical ground truth label for whether a genomic region originates from archaic introgression or not. We thus used existing introgression maps for these individuals as our introgression labels. Specifically, we used the introgression maps[29] inferred by the *HMMix* method[14] after filtering for introgression probabilities *>* 0.8. For the synthetically admixed testing individuals, we recovered the introgressed segments based on the *HMMix* introgression maps of the source individuals. We classified a window as being introgressed if over half of the window is covered by an *HMMix* tract.

#### 2.4.3 Testing data from the SGDP

We also evaluated the model’s performance when the testing data is sourced from a different dataset than the one used as a training dataset. To this end, we repeated the process for testing individuals in the *SGDP* dataset[25]. We take chromosome 1 of unadmixed individuals from three populations: West Eurasia *n* = 75, Africa without Yoruba *n* = 46, and East Asia *n* = 47. We again perform a forward-in-time admixture simulation using *haptools*[28], a human recombination map of chromosome 1[27], and simulation parameters of 25 generations of admixture with initial proportions Africa= ⅙, West Eurasia= 3/6, East Asia= 2/6. We use Yoruban individuals (of which there are only 3 in the *SGDP*) for *S\** calculations, and the AAFs of the unadmixed individuals in these three populations. In total, we simulate 200 admixed testing individuals from the *SGDP* data.

### 2.5 Model Validation

To validate our joint inference model, we measure inference accuracy under multiple scenarios (see Section 3.1), with varying training and testing datasets. We consider five scenarios:

i. Training and testing both on synthetic datasets.
ii. Training on the synthetic dataset with testing on the *1KG* dataset.
iii. Training on the *1KG* chromosome 1 dataset combined with the synthetic dataset, and testing on the *1KG* chromosome 1 dataset.
iv. Training on the *1KG* chromosome 1 dataset combined with the synthetic dataset, and testing on the *1KG* chromosome 22 dataset.
v. Training on the *1KG* chromosome 1 dataset combined with the synthetic dataset, and testing on the *SGDP* chromosome 1 dataset.

We also perform an ablation study to quantify the contribution of *S\** features and joint multitask training to both continental ancestry and archaic introgression inference. Starting from scenario **i**, which uses the full model with SNP embeddings *E_W_*, *S\** summary features, and both the continental ancestry and archaic introgression classifiers, we evaluate all possible ablations of these components. We consider a multi-task version of the model without *S\** information, and single-task versions of the model both with and without *S\** information. SNP embeddings *E_W_* are always included. This allows us to separately measure the effects of *S\** features and the benefit of joint training for each inference task. We denote these extensions of scenario **i** as **i-a)** through **i-e)**. Ablation results are reported in Table 1, where they can be directly compared to the full-model accuracies from scenario **i**.

**Table 1:**
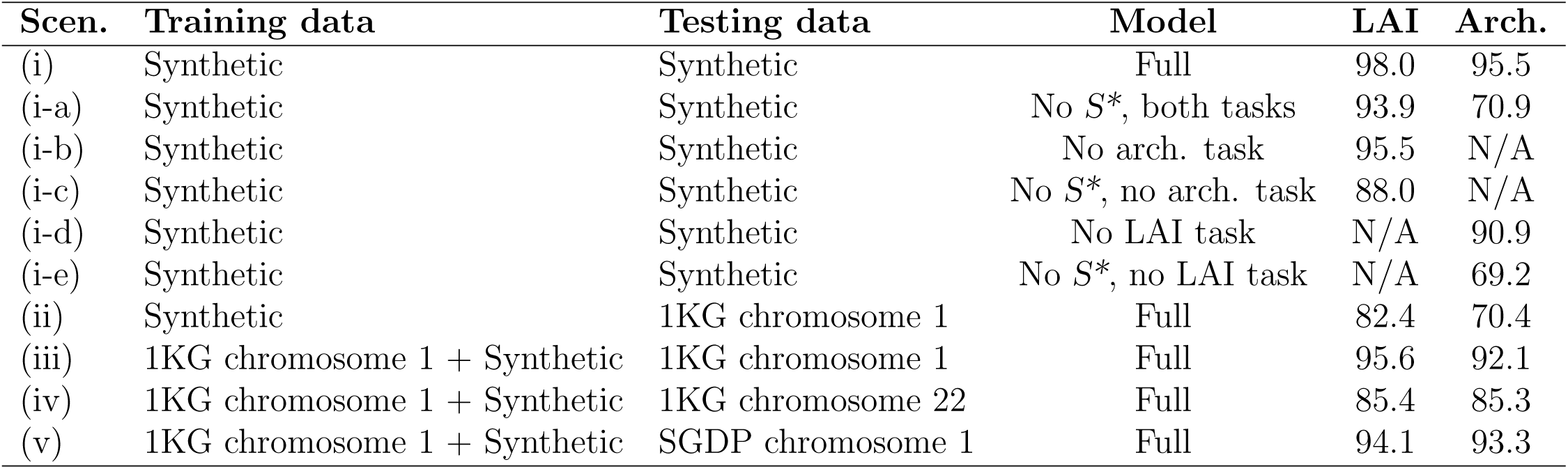
Model performance across multiple scenarios. For each scenario, we report continental local ancestry inference (LAI) accuracy and archaic introgression classification accuracy. **i)** Training and testing on synthetic data using the full model. **i-a)** through **i-e)** extend scenario **i)** with all non-empty combinations of the ablated *S\** input, continental ancestry classifier sub-network, and archaic introgression classifier sub-network. **i-a)** removes *S\** while retaining joint LAI and archaic introgression training. **i-b)** removes the archaic classifier sub-network. **i-c)** removes both *S\** and the archaic classifier sub-network. **i-d)** removes the continental classifier sub-network. **i-e)** removes both *S\** and the continental classifier sub-network. **ii)** Training on synthetic data and testing on *1KG*. **iii)** Training on *1KG* chromosome 1 + synthetic dataset and testing on *1KG* chromosome 1. **iv)** Training on *1KG* chromosome 1 + synthetic dataset and testing on *1KG* chromosome 22. **v)** Training on *1KG* chromosome 1 + synthetic dataset and testing on *SGDP* chromosome 1.

When reporting accuracy, we do it independently for continental and archaic ancestry. For continental ancestry inference accuracy, we report standard multi-class accuracy. Having the total number of windows *N*, and *y_i_, y*^*_i_* the true and predicted continental ancestry labels for window *i*, continental ancestry accuracy is calculated as:

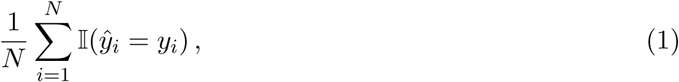

where I(*·*) is the indicator function.
For archaic introgression inference, we compute balanced accuracy, defined as the mean of the true positive rate and true negative rate, to account for class imbalance between introgressed and non-introgressed windows. After calculating the number of true positives *TP*, true negatives *TN*, false positives *FP*, and false negatives *FN*, balanced accuracy is calculated as:

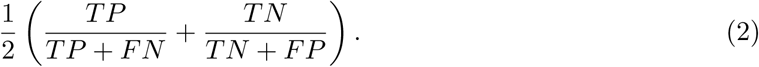

When measuring the accuracy of archaic inferences on real individuals, we again used the introgression maps[29] inferred by the *HMMix* method[14] as our ground truth.

### 2.6 Inference on admixed populations from the 1KG

We perform joint ancestry and introgression inference on 4 admixed populations from the *1KG*[30]: PUR *n* = 104, CLM *n* = 94, MXL *n* = 64, and admixed PEL *n* = 52. These individuals have genetic ancestry mainly from African populations, European populations that have been shown to be closest to Spanish populations[17][16], and Indigenous American populations. To infer the joint ancestry of these individuals, we again trained the model on a combination of real and synthetic data. For our real data, we trained on three populations representing African (AFR), European (EUR), and Indigenous American (AMR) genetic ancestries: Yoruba (YRI, *n* = 188), Iberians (IBS, *n* = 162), and unadmixed Peruvians (PEL *n* = 33). The 33 PEL individuals have been found by Reynolds et al. (2019)[31] to harbor almost no European or African ancestry. Here, YRI are also the outgroup population required to compute *S\** (Section 2.1). We specifically chose IBS as the population that most closely matches the European donor population during the time of American colonization. As previously described in the preceding subsections, we use introgression maps[29] inferred by the *HMMix* method[14] to generate our training labels for archaic or non-archaic windows.

We re-trained the model using 10,000 random, non-overlapping 1,000 SNPs windows from all available autosomes for these unadmixed individuals. Finally, these 10,000 windows are supplemented with the synthetic training data generated with demographic models *A, B* and *C* as described in Section 2.4, resulting in a total of 20,000 training windows encompassing both real and synthetic individuals. The order of these synthetic and real windows is randomized before being fed into the model for training.

### 2.7 Archaic ancestry sharing

When jointly inferring continental and archaic introgression of admixed populations in the *1KG* dataset, we analyzed the percentage of the archaic genome that we can recover from introgressed individuals within an Indigenous American or European genetic background; this allows us to test if there are any differences in archaic variation for different continental sources in admixed populations. We first identified all genomic windows where at least one introgressed haplotype was found. The fraction of the archaic genome recovered is the sum of the lengths of all the windows divided by the total physical size of the entire genome (all 22 autosomes). In addition, we keep track of the continental ancestry of these introgressed genomic regions. There are three possible cases: an introgressed haplotype can be found only on an Indigenous American background, only a European background, or found in both. These three scenarios let us calculate the percentage of the archaic genome recovered from archaic variation private to each continental ancestry, or shared between them. We also keep track of the proportion of introgressed windows for each continental ancestry, which gives us a measure of introgression density conditioned on continental ancestry.

Given that the admixed populations considered here have different levels of African, European and Indigenous American ancestry, and sample sizes are different, the fraction of the introgressed genome recovered would be biased by the population’s global continental ancestry proportions and sample size. For example, if a certain window for a population consists of mostly European haplotypes, a higher introgression density will be measured for the European ancestry due to the ancestry imbalance. To account for this, for each window *W* we balance the ancestry groups by sampling *n_w_*haplotypes without replacement:

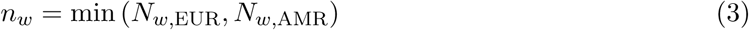

After sampling for each window, we further classify windows where at least one sampled haplotype is introgressed into three categories: introgression private to EUR haplotypes, private to AMR haplotypes, or shared by at least two haplotypes of different continental ancestries. The fraction of the whole genome assigned to category *p ∈* EUR-private, AMR-private, shared) is: Identifying candidates for adaptive introgression

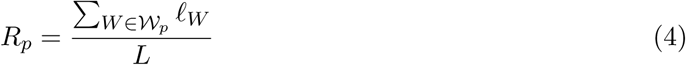

where *ℓ_W_*is the physical length of window *W*, *L* is the total physical length of the genome, and *W_p_* is the set of windows belonging to category *p*.

### 2.8 Local ancestry heterozygosity analysis

Given the continental ancestry calls for all individuals, we measured the per-population proportion of different ancestry pairings, considering the two haplotypes that make up a diploid individual. For any given window, an individual can have heterozygous continental ancestry (*e. g.*, AFR/EUR ancestry calls), or homozygous continental ancestry (*e. g.*, AMR/AMR ancestry calls). Furthermore, given the global continental and archaic ancestry proportions for a population, we can calculate the expectation for how frequent all possible ancestry pairs would be genome-wide, as well as the expectation for many of these pairs would harbor archaic introgression.

To simulate the null distribution of ancestry pairing within each population, we randomly sample 10,000 pairs of values from the possible continental ancestries: [*AFR, EUR, AMR*], the sampling probability for each ancestry is the same as the genome-wide proportion of that ancestry at the population level. For example, given the global continental ancestry proportions of the PUR population of *d* = [*∼* 0.15*, ∼* 0.2*, ∼* 0.65] for [*AFR, EUR,* and *AMR*], we construct 10,000 parings by randomly sampling from the 3 possible ancestries using *d* as the draw weights. Analogously, we sample for introgression at the pair level by using the population-level proportions of introgression within each continental ancestry. Having constructed these 10,000 ancestry pairs, we measure the proportion for each continental ancestry pairing. We repeat this process 100 times, thus finding 100 observations for randomly sampled proportions for all ancestry pairings, as well as 100 observations for frequency of introgression conditioned on continental ancestry pairs. We can then use the mean and standard deviation from these observations in addition to the actual observed proportions to conduct a Z-test, informing us on any statistically significant deviations from the expected ancestry pair proportions, or expected introgression within each pair category.

### 2.9 Identifying candidates for adaptive introgression

For each population, we identify candidate regions for adaptive introgression by combining estimates of introgression proportion with evidence of positive selection from integrated haplotype scores[32] (iHS). We first select genomic windows that satisfy two criteria: (i) at least 10% of haplotypes within the window were inferred to be introgressed, and (ii) the window either overlapped a gene or was located within 10 kb of a gene. For each qualifying window, we extend the window to a larger 50kb region centered around the qualifying window. In this 50kb region, we identify all SNPs with extreme standardized iHS values, referred to as critical SNPs (i.e. normalized *|*iHS*| >* 2; details of the iHS calculations are provided in Section S4 of the Supplementary Material). These 50kb windows represent candidate introgressed regions that also show evidence of recent positive selection.

If we identify critical SNPs in the 50kb window, we further extend the window by 5kb on each side to check if there are more critical SNPs nearby. If there are critical SNPs within 5kb of the edges of the region, the region is iteratively expanded by 5kb to capture all critical SNPs in close proximity. Finally, we retain a region as a candidate for adaptive introgression if at least 10% of the critical SNPs also carry an archaic allele (i.e., an allele present in a Neanderthal or Denisovan genome). These variants are referred to as archaic critical SNPs.

Having identified regions that pass all filters for being candidates of adaptive introgression, we sort the regions by the mean iHS value of the archaic critical SNPs within the region. For each population, we then select the top 20 regions and their associated genes. To build the final list of candidate genes for adaptive introgression, we take the union of these 80 genes, resulting in a final list of 63 genes (as some genes pass all filters in more than one population). For all the top genes, we report the mean absolute normalized iHS value for the archaic critical SNPs in the window overlapping the gene, introgression proportion, and continental ancestry background of introgressed tracts for each of the four admixed Latin American populations in the *1KG*. We note that even if a candidate gene was selected because of a high iHS value and a high introgression proportion in one population, it might have a low mean iHS value or introgression proportion value in another population.

## 3 Results

### 3.1 Training on both real and synthetic data improves multi-task accuracy

We detail the model accuracy after training and testing on multiple scenarios (Table 1) in Section 2.5. Overall, we find that the model achieves very high accuracies for both the continental (98.0%) and archaic inference (95.5%) when training and testing on completely synthetic data (Scenario i, Table 1). We find that training single-task versions of the model (Scenarios **i-b** and **i-d**, Table 1) reduces accuracies for both continental ancestry (94.4%) and archaic ancestry (90.9%) tasks when compared to the full multi-task model. This shows that model generality and performance for both tasks improves by joint training. We also find that removing *S\** feature information (Scenarios **i-a**, **i-c** and **i-e**, Table 1) results in reduced performance not just for the archaic introgression detection task, but also for the continental ancestry accuracy.

We also achieve accuracies considerably better than chance (82.4% continental, 70.4% archaic) when the model is trained on purely synthetic data and then applied to real human genomes (Scenario ii, Table 1). However, the best accuracies in real individuals are achieved when combining both real and synthetic datasets during model training. For example, after training on real genomic windows from chromosome 1 of the *1KG* in addition to synthetic windows, we achieve very high accuracies (95.6% continental, 92.1% archaic) on chromosome 1 of the *1KG* test individuals (Scenario iii, Table 1). When tested on a different chromosome (22) of the *1KG* individuals (Scenario iv, Table 1), the model also achieves high accuracies (85.40% continental, 85.3% archaic). The reason is that including AAF information in the input matrix provides information about the allele frequency distribution that is likely conserved across chromosomes. However, the accuracy is lower than the accuracy for chromosome 1, suggesting that training exclusively on chromosome 1 does not fully capture some features of chromosome 22.

We also achieve high accuracies even when we test the model on chromosome 1 of individuals that belong to a completely different sequencing project, namely the *SGDP* (Scenario v, Table 1, 94.1% continental, 93.3% archaic). We note that despite differences in the set of SNPs between the *SGDP* and the *1KG* data, the accuracy is almost identical to the accuracy achieved when the testing genomes come from the same sequencing project as those used for training. Overall, our benchmarking indicates that our method accurately infers both continental and archaic ancestry tracts and that jointly modeling these two sources of ancestry yields higher accuracy than inferring either in isolation. Importantly, we demonstrate that we can combine synthetic and real genetic data to achieve the highest accuracy, and we can perform cross-dataset inference on real human genomes. These results highlight the model’s capacity to generalize across heterogeneous datasets.

### 3.2 Indigenous American tracts show the highest proportion of archaic introgression

We applied our method to 4 admixed populations in the *1KG*: PUR (*n* = 104) Puerto Ricans from Puerto Rico, CLM (*n* = 94) Colombians from Medellin, Colombia, MXL (*n* = 64) Mexican Ancestry from Los Angeles, and admixed PEL (*n* = 52) Peruvians from Lima, Peru. In total, that is 314 admixed individuals in 4 populations. We retrained our model using the IBS population as the European source because Latin Americans derive their European ancestry from mostly Spanish ancestors. We used unadmixed PEL individuals as the reference for the Indigenous American genetic ancestry (see section 2.6 for more details). We inferred both continental and archaic ancestry tracts along the genome for all haploid genomes in MXL, CLM, PUR and PEL samples. After inferring the continental and archaic ancestry tracts for all individuals in each population, we computed the continental ancestry and introgression proportion distributions for each haploid genome in our sampled individuals (Figure 2). For the global continental ancestry proportions, our results (Figure 2, panel A) are comparable to previous studies on these populations[17][16]. For example, PEL shows the highest proportion of Indigenous American ancestry, even when considering only the admixed PEL individuals[31], with an average Indigenous American proportion of *∼* 60%. Notably, we infer slightly higher average Indigenous American proportions for PUR and CLM individuals (*∼* 20% and *∼* 30%, respectively) than previously reported[17][16] figures (*∼* 13% for PUR and *∼* 26% for CLM). This is particularly significant for the PUR population, as we find that, on average, Indigenous American ancestry is more common than African ancestry for PUR individuals, while previous studies find the opposite[17][16]. This could be related to the use of a different reference panel of Indigenous American individuals used by these studies[33]. Levels of European ancestry are highest in PUR and CLM, and in MXL the distribution has a larger variance (Figure 2, panel A). The proportion of African ancestry is highest in CLM and PUR but the proportion of African ancestry is on average smaller than the proportion of European ancestry. Measuring introgression per individual shows that PEL individuals exhibit the highest proportion of introgression, followed by MXL, CLM and PUR. Similarly to Witt et al.[34], we are able to reproduce the observation that archaic ancestry is positively correlated with Indigenous American ancestry using genome-wide proportions (see Supplementary Figure S1). Notably, here we can investigate the relationship more deeply because we have joint calls of both continental and archaic ancestry along the genome. Consequently, we computed the proportion of introgressed tracts within each continental ancestry per haploid genome to obtain a distribution (Figure 2, panel B). As expected, the proportion of archaically introgressed AFR tracts is close to 0% for all populations. We also find that, for the CLM, PEL, and MXL populations, there is a higher likelihood for Indigenous American tracts to carry introgression than European tracts. On average, we find that *∼* 1.7% of European tracts contain introgression, and *∼* 2.0% of American tracts show contain introgression. These results are consistent with the global introgression proportions (Figure 2, panel C), as the highest mean introgression proportion (*∼* 1.7%) is observed in PEL, which has the highest proportion of Indigenous American ancestry at *∼* 60%. We find that the main factor of overall introgression proportion is the amount of Indigenous American ancestry found in a population (see the linear regression analysis in Supplemental Figure S1; *R*^2^ = 0.562, *p* = 1.88 *×* 10*^−^*^114^).

**Figure 2:**
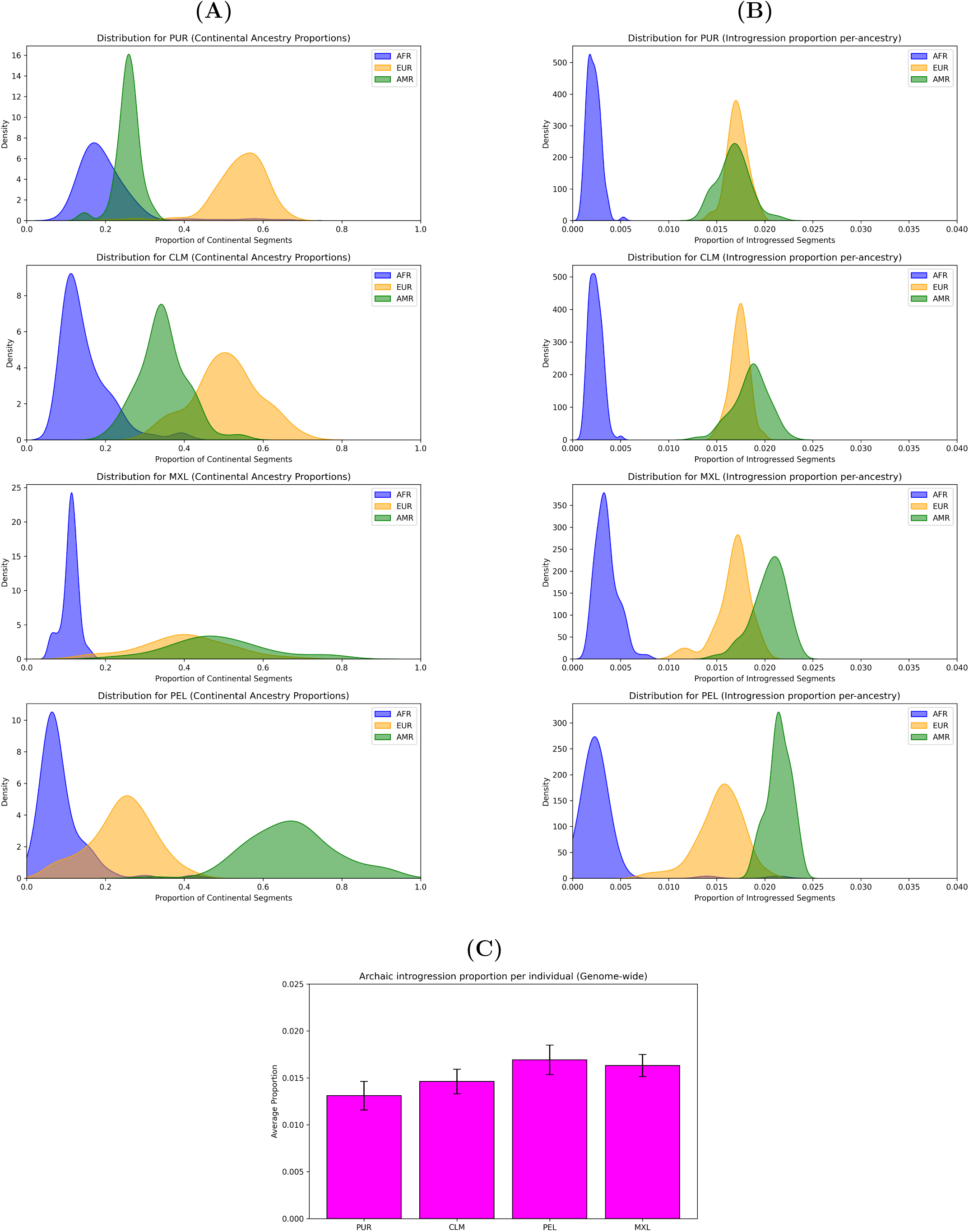
Global ancestry proportions are calculated on a per-haploid genome basis to create distributions. **(A)** Genome-wide distributions of proportion of AFR, EUR, and AMR genetic ancestries. **(B)** Genome-wide distributions of proportion of introgressed tracts conditioned on continental ancestry. **(C)** The mean genome-wide introgression proportion per population; error bars indicate standard deviation.

### 3.3 Indigenous American and European tracts contribute different fractions of recoverable archaic sequences

We constructed 100 unadmixed Indigenous American and European pseudo-genomes through sampling of the admixed individuals in each population, as described in Section 2.7 (Figure 3). We then used these pseudo-genomes to measure how archaic introgression changes in proportion and distribution through the genome, conditioned on continental ancestry. Even though Indigenous American ancestry is more likely to carry archaic ancestry (see Section 3.2), we recover a higher fraction the archaic genome (measured as the fraction of the genome where at least one pseudo-genome shows signs of introgression) from European tracts (Figure 3, panel A). On average, we recover *∼* 7% of the genome through archaic tracts private to American tracts, *∼* 14% of the archaic genome from archaic variation private to European tracts, and *∼* 6% of the genome from introgressed tracts with a mixed continental background.

**Figure 3:**
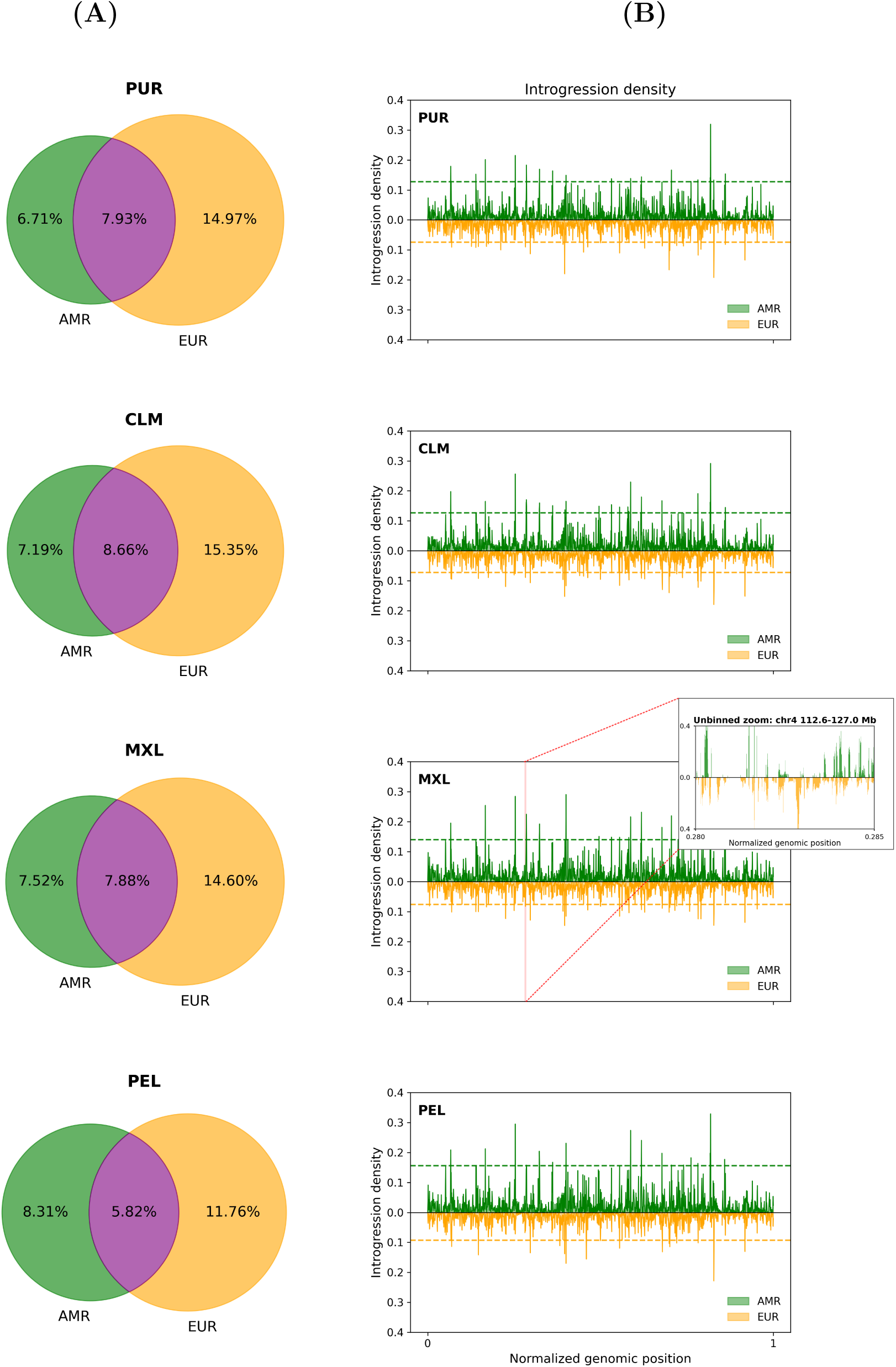
**(A)** Percentage of genome recovered from: only Indigenous American tracts (green), only European tracts (orange), or shared by both continental ancestries (purple). **(B)** Proportion (y-axis) of introgressed Indigenous American (green) or European (orange) haplotypes for a given genomic position (x-axis), a dashed line indicates the mean proportion of haplotypes showing evidence of introgression in genomic regions where at least one haplotype is introgressed. **(Insert)** Throughout the genome, introgression is more diffusely distributed across European ancestry windows.

We find that introgression in Indigenous American tracts tends to be shared by many haplotypes, resulting in a higher average introgression proportion and a lower fraction of the genome covered. To illustrate this, we calculated the proportion of sampled haplotypes that show evidence of archaic introgression at each genomic position (Figure 3, panel B), which gives us a measure of archaic ancestry density in both Indigenous American and European backgrounds. We also calculated the average introgression proportion at the population level, conditioned on continental ancestry, for regions where at least one haplotype is introgressed (Figure 3, panel B, dashed lines). We find that in genomic windows where at least one Indigenous American haplotype is introgressed, the average proportion of introgressed Indigenous American haplotypes is *∼* 13%. If we instead condition on European haplotypes, the average proportion drops to *∼* 8%. These results suggest a higher degree of relatedness among the sampled Indigenous American haplotypes. This behavior is the same within each of the studied populations.

### 3.4 Ancestry heterozygosity deviates from random expectation

For every genomic window in each population, we keep track of every diploid individual’s ancestry pairing. Individuals can either be homozygous (*e.g.*, *AFR/AFR*) or heterozygous (*e.g.*, *EUR/AMR*) for a given window. At the same time, an individual can show evidence of archaic introgression in a genomic window if at least one haplotype harbors archaic ancestry. Given the global continental and archaic ancestry proportions for each population (Figure 2, panel A), we can also calculate the expected distributions of each ancestry pairing and their introgression proportion as described in Section 2.8. Figure 4 shows the observed and expected proportion of continental ancestry pairings and archaic introgression proportion for each population. We find that in the four populations, homozygous ancestry pairings are over-represented compared to a baseline expectation, while heterozygous pairings are under-represented, this discrepancy is statistically significant in all cases. This over-representation of ancestry homozygosity can be consistent with assortative mating in post-conquest Latin America[18], or indicative of population substructure within these admixed populations (*e.g.*, regional population structure in Colombia[35]). When looking at the expected and observed archaic introgression proportions for the different continental ancestry pair categories, we do not find any notable deviations from expectation. However, we observe that the highest proportion of introgression is consistently found in the *EUR/AMR* pairs. This is expected, as we are effectively combining two sources of archaic introgression.

**Figure 4:**
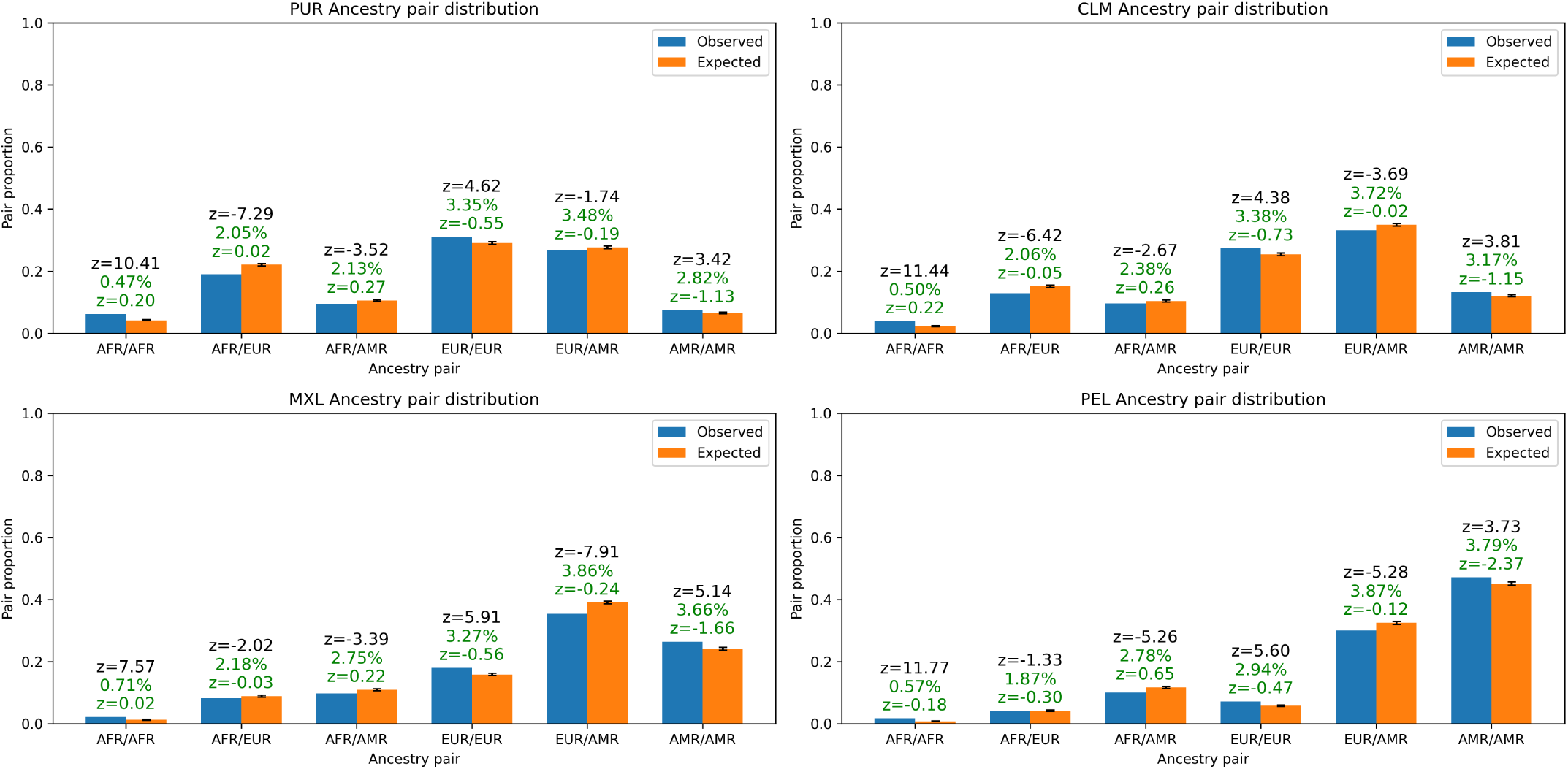
Observed (blue) vs. expected (orange) ancestry pair proportions for each studied population. We find that in all populations, homozygous ancestry pairings are significantly more common than expected by chance. Z-test scores (in black) for continental ancestry proportions are shown above each pair category. Error bars on the orange bars indicate the standard deviation over 100 observations. Percentages (in green) and Z-test scores (in green) show the observed proportion of introgression within each continental ancestry pair category as well as deviation from expectation. We do not find that observed introgression proportion within each category significantly deviates from expectations.

### 3.5 Continental ancestry background of putatively adaptive introgressed loci

An advantage of jointly inferring archaic and continental ancestry is that it allows us to characterize the continental ancestry context of putatively adaptive introgressed haplotypes. This allows us to determine whether these introgressed segments were inherited through Indigenous American or European ancestral lineages. To identify candidates for adaptive introgression, we obtain iHS scores for MXL, PEL, PUR and CLM (see Section 2.9) individuals. For each population, we then identify the genomic regions with at least 10% introgression proportion, and within this set, we find the the top 20 genes with the highest iHS values. Some genes are identified in multiple populations, combining them leads to a total of 63 distinct genes with signatures of both introgression and positive selection. Since we also have calls for continental ancestry, we also query whether the putative selected introgressed segment is within an Indigenous American or European genetic background. Figure 5 shows the values for iHS (panel A), the introgression proportion (panel B), and the continental ancestry composition (panel C) of introgressed tracts overlapping these 63 genes in all four populations.

**Figure 5:**
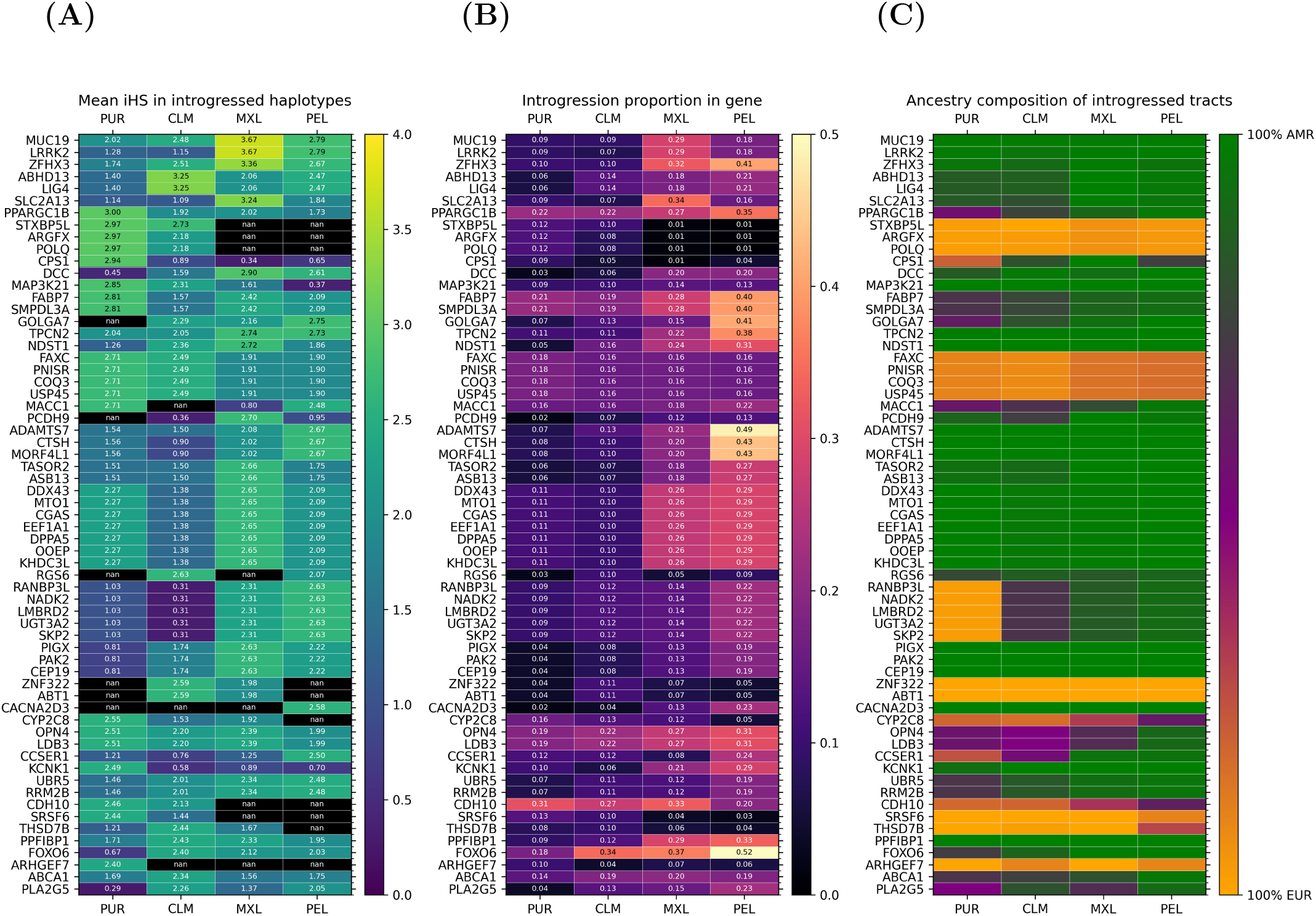
**(A)** Mean absolute iHS calculated within introgressed haplotypes for the top candidate genes. **(B)** Proportion of individuals per-population that show evidence of archaic introgression in regions overlapping the candidate genes. **(C)** Continental ancestry composition of introgressed tracts overlapping each gene. Green cells indicate that all introgressed tracts in the region have a Indigenous American ancestry, orange corresponds to European ancestry, and purple to an even mix of continental ancestries.

We find that most candidate genes have a maximum iHS *>* 2.5, which is the threshold for the most extreme 1% of iHS values genome-wide[32] (Figure 5, panel A). We find 7 candidate genes (*MUC19*, *LRRK2*, *ZFHX3*, *ABHD13*, *LIG14*, *SLC2A13*, *PPARGC1B*) with maximum iHS *≥* 3.0, indicating even more extreme haplotype homozygosity signals, for these 7 genes, the archaic haplotype is practically always contained within an Indigenous American continental ancestry (Figure 5, panel C). Moreover, we find that introgression proportion in these 7 genes is always high (*>* 20%) in either the PEL or MXL populations, consistent with elevated Indigenous American ancestry (Section 2). To our knowledge, the *ABHD13*, *SLC2A13*, and *PPARGC1B* genes have not previously been described as candidates for adaptive introgression.

Some of the genes found by this approach have already been identified as likely targets of adaptive introgression, such as *MUC19*[21] and *ZFHX3*[1]. Remarkably, we find that 41% of PEL individuals show evidence of introgression in the *ZFHX3* region (Figure 5, panel B), looking at all candidate genes, we find eight instances where the introgression proportion for PEL individuals is *>* 40% (Figure 5, panel B). In general, the introgressed tracts that overlap a given gene also tend to belong to the same continental ancestry, with Indigenous American ancestry being the most common (Figure 5, panel C). This suggests that most candidates for adaptive introgression in admixed populations of Latin America correspond to selective events that may be older than European colonization. However, selection in candidate genes with a mixed continental background (*e.g. OPN4*) could be consistent with post-colonial selection of archaic variants present in both European and Indigenous American ancestries.

## 4 Discussion

In this work, we present the first deep-learning method for the joint inference of continental and archaic ancestry (Figure 1). We find that by combining SNP sequences, AAF information, and summary statistics that capture signals of archaic introgression such as *S\**, we are able to accurately infer both continental and archaic ancestry. For example, we show that supplementing SNP sequence data with statistics typically reserved for archaic ancestry prediction improves the accuracy of our continental ancestry calls (Table 1). The reverse holds true as well, with our archaic ancestry predictions achieving higher accuracies; likely because the continental ancestry signal present in the SNP embedding also informs about *S\** score significance thresholds (Table 1). We show that models trained exclusively on synthetic data achieve strong performance when applied to real genomes (*e.g.*, the 1KG dataset), reaching accuracies of *∼* 82% for continental ancestry and *∼* 70% for archaic ancestry. While simulated data has traditionally been used to infer demographic parameters through summary statistics, our results demonstrate that demographic models inferred in this way can in turn generate synthetic genomes that capture key features of real genetic variation and enable accurate evolutionary inference along the genome.

Our approach partly relies on the capability of simulating genomic datasets that realistically replicate some features of real datasets[24], which has been the common way that machine learning methods are trained in the field of population genomics[36][37][38]. However, we achieve the highest accuracy when we train on both synthetic and real data; using synthetic datasets as a way to augment the training dataset (Table 1, scenario iii). This strategy has previously been shown useful for biological tasks where data scarcity is an issue[39], as well as specifically for continental ancestry inference tasks[40]. Finally, this hybrid training mitigates the “reality gap” inherent to simulation-based frameworks by encouraging the model to perform well on both synthetic and real data. It improves robustness to domain shift, which can occur due to differences in genotyped SNPs between the training and testing individuals (cross-dataset) or by genetic drift between the admixed populations being analyzed and the reference panels treated as source populations (within-dataset); all this without explicitly learning an adaptation objective. Future work could expand on this approach by incorporating explicit domain adaptation techniques, which have been shown to improve model performance in population genetics tasks where labeled real data is unavailable[41], as is the case for archaic introgression in human datasets[42]. We further show that our model can be trained on individuals from one sequencing project (*e.g.*, the *1KG*), and then applied to individuals from a completely different dataset (*e.g.*, the *SGDP*). This is possible because of how our model is trained on the relationships between allele frequencies in the reference populations and the query haplotypes, which are combined into the model input (Section 2.3).

To demonstrate the utility of our model, we also apply our method to four admixed Latin American populations from the *1KG* (Figure 2). Our genome-wide measures of continental and archaic ancestry proportions are similar to those reported in previous studies[17][16], demonstrating the overall robustness of our results. One key advantage of our method is that we can quantify how often archaic segments co-occur with European or Indigenous American chromosomal segments, allowing us to infer whether an archaic tract was introduced via an Indigenous American or European ancestor. We find that, just as genome-wide continental genetic ancestry proportions vary across populations, the introgression proportion within a particular continental genetic ancestry varies between Latin American populations (Figure 2, panel B). We find that Indigenous American tracts are more likely to show evidence of introgression in MXL, PEL, and CLM individuals (*∼* 2.2% of tracts in MXL and PEL) than European tracts (*∼* 1.7%), and African tracts exhibit the lowest introgression proportion in all populations (*∼* 0.2%). This is likely due to Indigenous American tracts having a similar background level of Neanderthal introgression as European tracts[43], in addition to harboring some Denisovan ancestry introduced through introgression events that did not impact European populations[44]. Consequently, we find that the proportion of archaic ancestry in admixed Latin American populations is correlated with the total proportion of Indigenous American ancestry, consistent with previous studies[1] (Figure 2, panel C; Supplemental Figure 1).

We measure the fraction of the archaic genome recovered from a sample of introgressed Indigenous American and introgressed European tracts (Figure 3). In all populations, we find that a higher fraction of the archaic genome is recovered from segments of European genetic ancestry. This is true even when the global Indigenous American ancestry proportion is higher than the European proportion. The reason is that introgression is more evenly distributed among European tracts, while Indigenous American haplotypes have a higher likelihood of sharing archaic introgression in the same genomic region (Figure 3, panel B). This parallels previous observations in comparisons of European and East Asian populations: although East Asian harbor higher levels of archaic ancestry than Europeans at the individuals level, more of the archaic genome is recovered from an equal-sized sample of Europeans[34]. This may reflect the stronger effects of genetic drift in East Asian and Indigenous American populations, resulting in an increase in the level of shared archaic ancestry between individuals. Alternatively, it could point to having European donors from distinct geographical locations. Future studies inferring tracts of identity by descent (IBD) in admixed individuals[17] and partitioning by continental ancestry may reveal whether the genetic composition of European donors influences the amount of archaic ancestry recovered. It should also be noted that sampling strategies in the *1KG* can bias these results, as the studied individuals are not necessarily representative of all populations in these countries.

For each population, we quantify the amount of genome within every possible ancestry pair across all individuals (six pairings possible, see Figure 4), and we find a statistically significant overrepresentation of homozygous ancestry pairs (AFR/AFR, EUR/EUR or AMR/AMR), suggesting population substructure[35] or possibly assortative mating[18]. Previous studies have identified assortative mating correlated with different continental ancestry backgrounds in Latin American populations, specifically linked to social stratification[18], which may lead to an over-representation of homozygous ancestry. We find that the highest proportion of introgression (*∼* 3.5%, Figure 4) is within regions of EUR/AMR heterozygous continental ancestry. This is consistent with our results on the archaic sequence recovered from each continental ancestry (Section 3.3), since there is a considerable proportion of archaic variation private to both the Indigenous American and European ancestries, heterozygous pairs have a larger pool of potential archaic introgression.

Finally, we combine our joint ancestry calls with integrated haplotype scores (iHS) to find candidate loci for adaptive introgression (Figure 5). We found multiple genes with evidence of selection on archaic variants, including some genes that have been identified by previous studies (e.g. *MUC19* and *ZFHX3*) [21][1]. It should be noted that this approach for finding candidate regions is conservative, as increased heterozygosity and shorter haplotypes in admixed populations[45] will lead to lower overall iHS values. We found that, in most cases (Figure 5, panel C), the adaptive introgressed segments are within an Indigenous American background. This provides evidence that the timing of these selective events may be older than European colonization, and may have helped humans adapt to the new environments of the Americas. This may also be consistent with an extremely strong selective pressure on the Indigenous American haplotypes following European colonization. Future studies using ancient genomes from the Americas will provide useful information to infer the timing of positive selection.

There are multiple aspects of our method that we can improve further. For example, one of the main difficulties of training machine learning models for archaic ancestry inference is the lack of ground-truth labels for archaic tracts in real genomes. Our method partly relies on existing introgression maps made by other methods (such as *HMMix*[14]). However, we achieved archaic-ancestry accuracies of *∼* 70% when training exclusively on synthetic introgressed genomes, and this provides confidence of our method’s performance even when existing introgression maps are not perfect. Potential extensions that could improve the recall of archaic tracts include taking a consensus-vote of multiple archaic inference methods, or including archaic sequences themselves as part of the model input. Finally, the inclusion of archaic sequences as part of the model input or as part of a post-processing step could let us identify not just the presence of introgression, but also identify the archaic donor population. Currently, our method can be paired with sequence similarity workflows to assign donor populations to the archaically introgressed tracts[15][46].

In summary, we present the first deep learning framework for the joint inference of continental and archaic ancestry. Our approach accurately infers both recent modern human ancestry and ancient archaic introgression. By training on a combination of real and simulated data, the model generalizes across diverse genomic datasets without requiring retraining for each new dataset. Applied to admixed populations from the 1,000 Genomes Project, our method provides a detailed characterization of the distribution of archaic ancestry and enables ancestry-specific analyses that were previously difficult to perform. In particular, it allows the continental ancestry of adaptive introgression candidates to be inferred, addressing an important gap in studies of adaptive introgression in Latin American populations[47].

Our analyses reveal that historical demographic processes have shaped the distribution of archaic ancestry differently across admixed populations, reflecting each population’s unique admixture history and intensity. We also identify candidate regions of adaptive introgression and, by resolving their continental ancestry, propose that many represent cases in which selection acted on standing archaic variation after modern humans expanded into the Americas. This hypothesis can be tested as additional ancient genomes from the Americas become available. More broadly, our findings suggest that although many examples of adaptive introgression reflect rapid adaptation following introgression[48], some archaic alleles persisted as standing genetic variation within modern human populations before later contributing to adaptation as humans expanded into new environments[49]. Applying our framework to larger collections of admixed genomes will enable increasingly powerful ancestry-specific analyses and provide deeper insight into the evolutionary impact of archaic introgression.

