## Supplemental information for "Joint ancestry inference reveals the landscape of archaic introgression in admixed populations"

August 29, 2026

**Jazeps Medina Tretmanis**[1], **Valeria Añorve-Garibay**[1], **David Peede**[2], **Mayra M. Bañuelos**[1,2]
**María C. Ávila-Arcos**[3]*, **Flora Jay**[4]*, **Emilia Huerta-Sanchez**[1,2,5]*

[1] Center for Computational Molecular Biology, Brown University.

[2] Department of Ecology, Evolution and Organismal Biology, Brown University.

[3] International Laboratory for Human Genome Research, UNAM.

[4] Interdisciplinary Laboratory of Numerical Sciences, Université Paris-Saclay.

[5] Data Science Institute, Brown University.

* Contributed equally.

### S1. Continental vs. archaic ancestry proportions

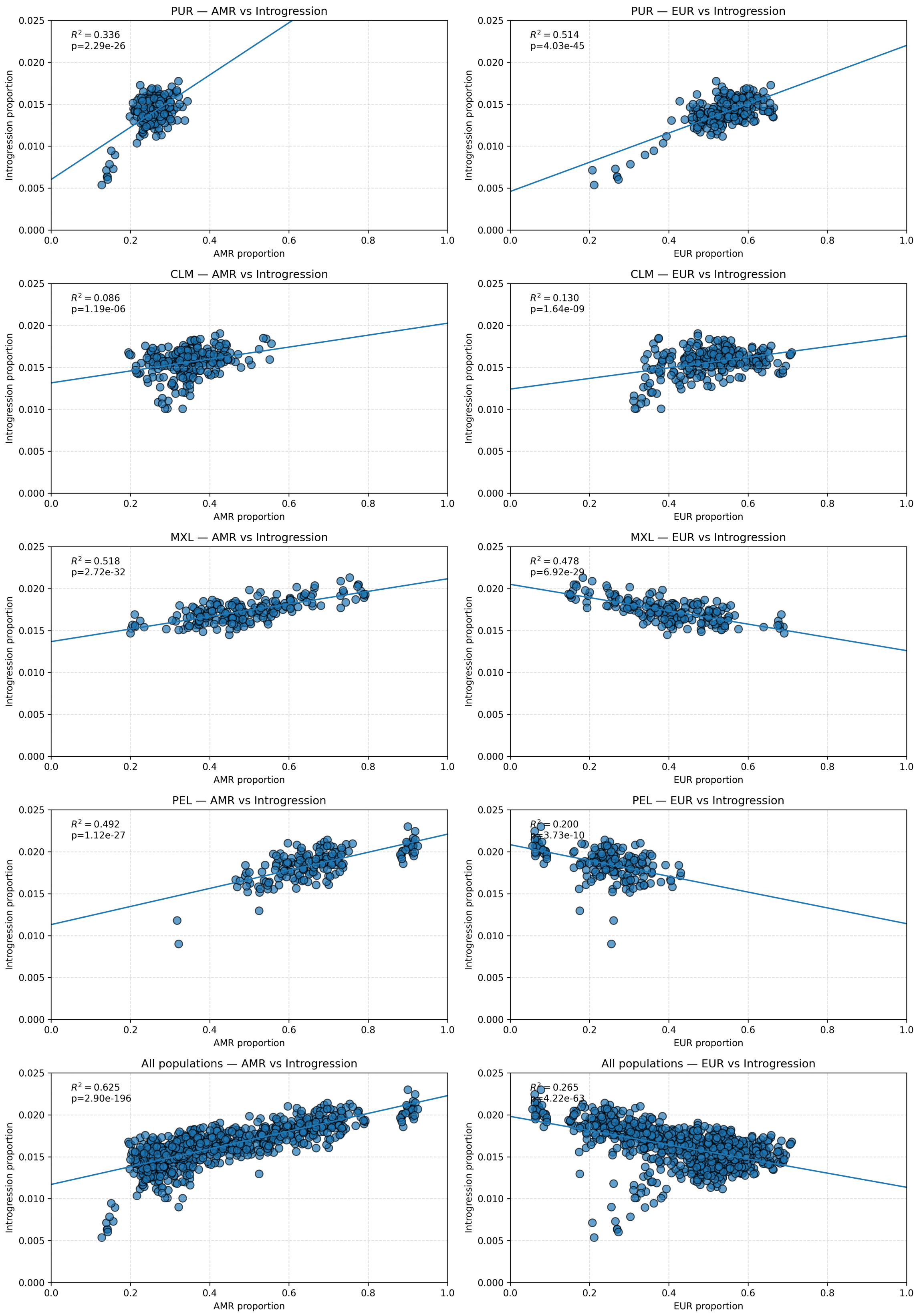


Figure 1: Native American (x-axis, left column) and European (x-axis, right column) proportion plotted against introgression proportion (y-axis). Each dot represents genome-wide proportions for a single haplotype. There is a clear, statistically significant positive correlation between Native American ancestry proportion and archaic introgression proportion

#### S2. CNN model layer description

The final model was selected through a hyperparameter grid search over learning rates $\{5 \times 10^{-5}, 5 \times 10^{-4}, 5 \times 10^{-3}\}$ and batch sizes $\{16, 32, 64\}$. The parameter combination that led to the lowest validation loss was chosen.

The continental ancestry (LAI) loss is calculated through a cross entropy loss function:

$$\mathcal{L}_{\text{LAI}} = \text{CrossEntropyLoss}(y_{\text{lai}}, \hat{y}_{\text{lai}}) \tag{1}$$

The archaic ancestry task loss is calculated through a weighted cross entropy loss. A positive class weighing factor of 7.0 is applied due to the strong negative example bias of archaic introgression.

$$\mathcal{L}_{\text{ARC}} = \text{BCEWithLogitsLoss}(\hat{y}_{\text{arc}}, y_{\text{arc}}) \tag{2}$$

We use the ADOPT[1] optimizer, a modification of the Adam optimization algorithm which guarantees convergence at an optimal rate for any $\beta_2$ optimizer parameter.

| **Layer** | **Type** | **Description** |
|---|---|---|
| L1 | Conv1d ($c + 1 \rightarrow 4$) | Initial feature extraction from SNP + AAF matrix with kernel size 3, stride 1, padding 1. |
| L2 | Conv1d (4→4) | kernel size 3, stride 1, padding 1. |
| L3 | Conv1d (4→8) | kernel size 3, stride 1, padding 1. |
| L4 | Conv1d (8→8) + AvgPool | kernel size 3, stride 1, padding 1. |
| L5–L6 | Conv1d (8→16) | kernel size 3, stride 1, padding 1. |
| Pool1 | AvgPool1d | – |
| L7–L8 | Conv1d (16→32) | kernel size 3, stride 1, padding 1. |
| Pool2 | AvgPool1d | – |
| L9–L10 | Conv1d (32→64) | kernel size 3, stride 1, padding 1. |
| Pool3 | AvgPool1d | – |
| L11–L12 | Conv1d (64→128) | kernel size 3, stride 1, padding 1. |
| Pool4 | AvgPool1d | – |
| L13–L15 | Conv1d (128→256) | kernel size 3, stride 1, padding 1. |
| Pool5 | AvgPool1d | – |
| L16–L18 | Conv1d (256→512) | kernel size 3, stride 1, padding 1. |
| Pool6 | AvgPool1d | – |
| L19–L21 | Conv1d (512→512) | kernel size 3, stride 1, padding 1. |
| Flatten | Reshape | Transforms CNN embeddings into vector for task-specific MLPs. The CNN embeddings are flattened into a vector of length $2,048$, and concatenated with the $S^*$ and private SNP distribution information, resulting in a vector $E_W$ of length $2,054$. |
| LAI Network | MLP (3 hidden layers) | Fully connected network with dropout (p=0.2), outputs 3 LAI predictions corresponding to AFR, EUR, and EAS ancestries. Input and hidden layer size: $2,054$. |
| ARC Network | MLP (3 hidden layers) | Identical to LAI MLP, with binary output corresponding to presence or non-presence of archaic introgression. Input and hidden layer size: $2,054$. |

Table 1: CNN architecture for joint LAI and archaic ancestry inference.

### S3. Demographic model description

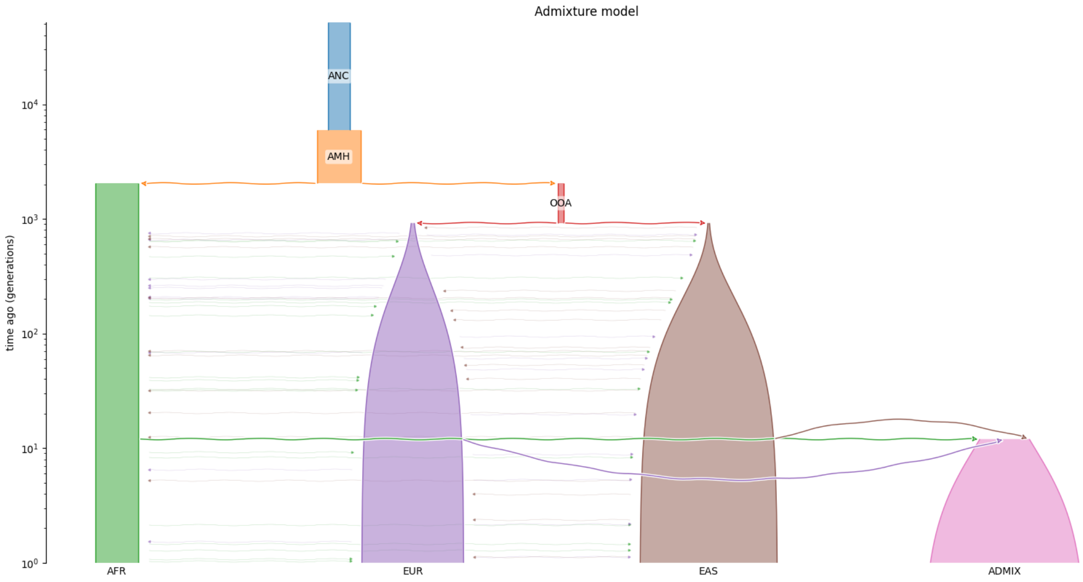


Figure 2: American admixture model with no introgression[2]. Simulated 2, 500 windows of 1, 000 SNPs each.

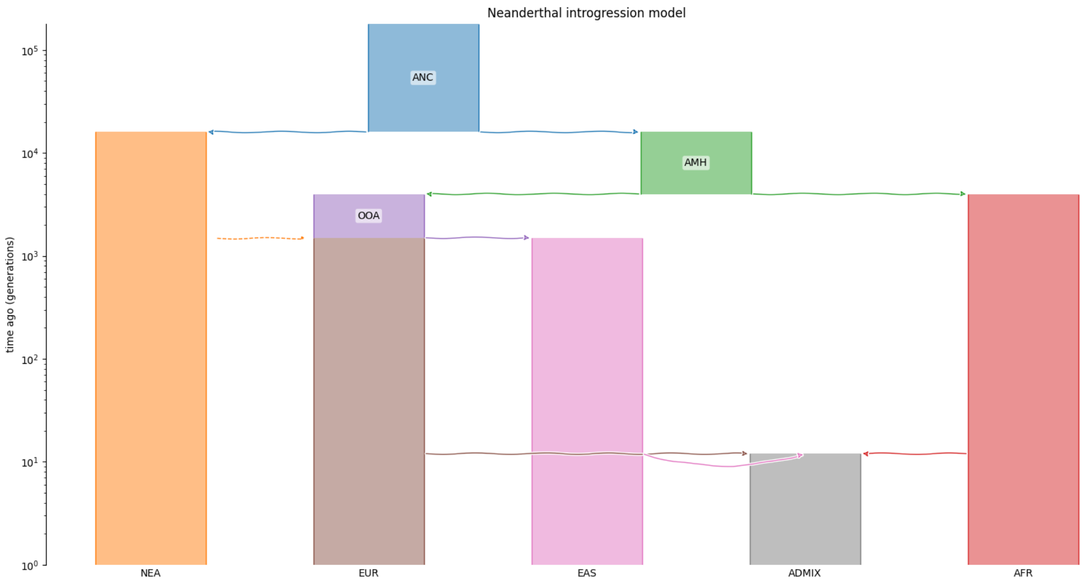


Figure 3: American admixture model with no introgression[2]. Simulated 5, 000 windows of 1, 000 SNPs each. 5 Replicates are simulated, with the only differences between replicates being the NEA introgression proportion (Values used: 1%, 2%, 5%, 10%, 15%).

Visualizations of demographic models done using the *demes*[4] and *demesdraw*[5] libraries.

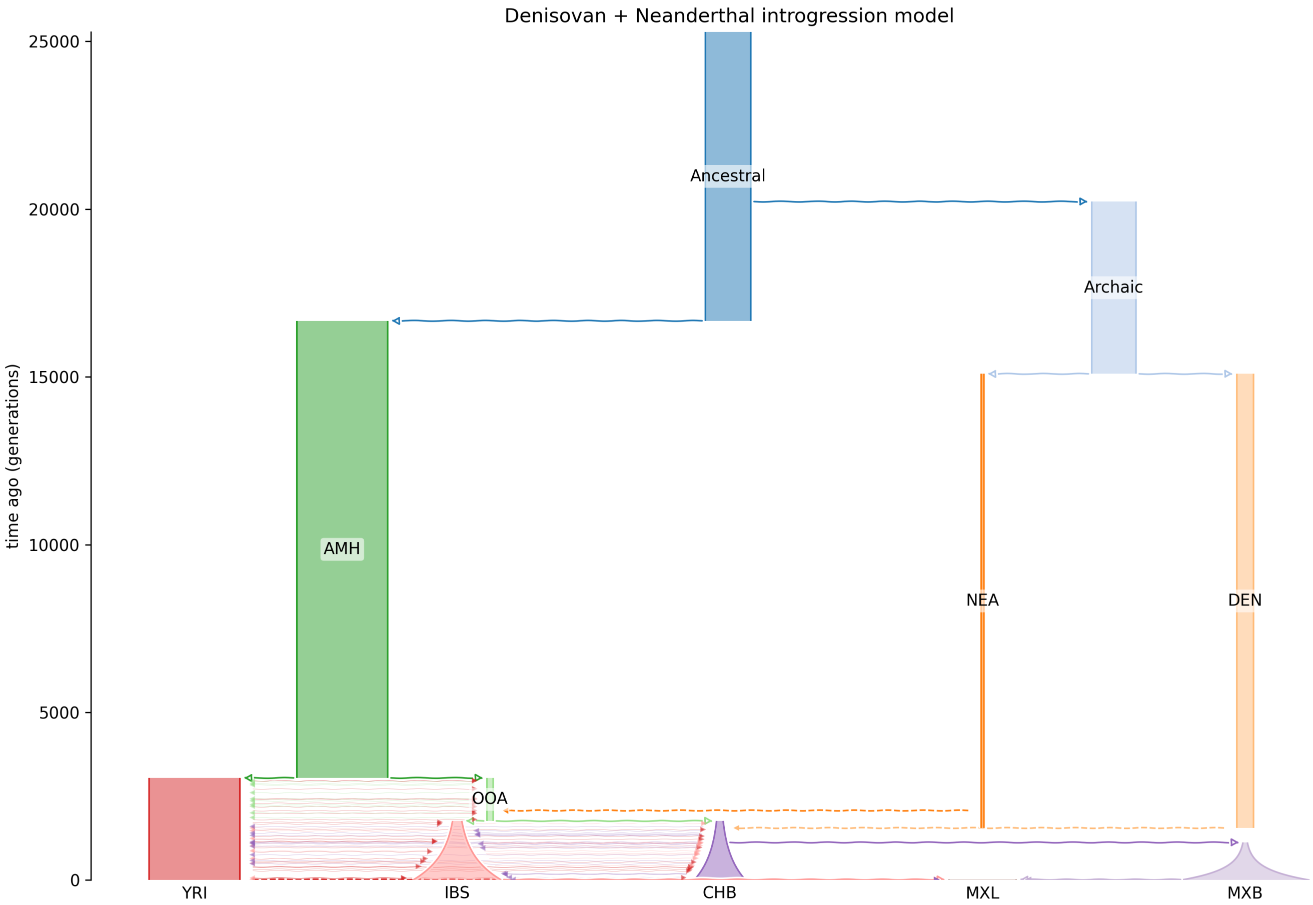


Figure 4: Neanderthal and Denisovan introgression model[3]. We simulated 2,500 windows of 1,000 SNPs each under a demographic model with anatomically modern human, Out-of-Africa (OoA), African (YRI), European (IBS), East Asian (CHB), Indigenous American (MXB), and admixed Mexican populations (MXL). The model includes one Neanderthal pulse from NEA into the OoA population, $p = 15\%$, at 2,069 generations ago, and one Denisovan pulse from DEN into the CHB, $p = 15\%$, at 1,552 generations ago. Mexican ancestry is modeled through MXL as an admixed population derived from MXB and IBS, with additional recent pulse contributions from YRI and CHB.

### S4. Algorithm: labeling introgressed tracts in simulated genomes

```
import numpy as np

def IntrogressedTracts(ts, target_populations, intro_populations,
   introgression_times):
    ret = []
    # Extract all information needed for computing
    # introgressed segments from migration
    migration_table = ts.dump_tables().migrations
    migration_nodes = migration_table.node
    migration_destinations = migration_table.dest
    migration_times = migration_table.time
    migration_left = migration_table.left
    migration_right = migration_table.right

    # Get archaic and modern samples
    # Returns samples in form of list even if single sample
    tar_haplotype = []
    for pop in target_populations:
        tar_haplotype.extend(ts.get_samples(pop))
    tar_haplotype = sorted(tar_haplotype)

    # Masks: Introgressing population and introgression time
    introgression_mask = np.isin(migration_destinations, intro_populations)
    within_time_mask = np.isin(migration_times[introgression_mask],
       introgression_times)

    # Get introgressing nodes for given introgression time and tree
       coordinates
    intro_nodes = migration_nodes[introgression_mask][within_time_mask]
    intro_left = migration_left[introgression_mask][within_time_mask]
    intro_right = migration_right[introgression_mask][within_time_mask]

    for tree in ts.trees(tracked_samples=tar_haplotype):
        # Set of leaf nodes in target population with introgressed tracts
```

```
        tree_intro = set()
        tar_set = set(tar_haplotype)

        for idx, node in enumerate(intro_nodes):
            if ((tree.interval[0] <= intro_right[idx] and intro_left[idx] <=
                tree.interval[1])):
                leaves = set(tree.get_leaves(node))

                if tree.num_tracked_samples(node) > 0:
                    to_remove = []
                    for target in tar_set:
                        if target in leaves:
                            tree_intro.add(target)
                            to_remove.append(target)
                    for x in to_remove:
                        tar_set.remove(x)

        sample_intro_dict = {}
        for sample in tar_haplotype:
            if sample in tree_intro:
                sample_intro_dict[sample] = 1
            else:
                sample_intro_dict[sample] = 0

        ret.append((tree.interval.left, tree.interval.right,
            sample_intro_dict))

    return ret
```

### S5. $S^*$ calculations

In addition to calculating the AAFs for a genomic window $W$, we also calculate its $S^*$ score distribution. For each window $W$, $S^*$ scores and the number of SNPs absent in the outgroup population (we used Yoruban individuals in the *1KG*) are calculated in sliding windows of 50kb[6] with a step size of 10kb. For each of these sliding windows, we also record the number of SNPs not present in the outgroup population (specifically, Yoruban individuals in the *1KG*) that are associated with each $S^*$ score. We note that while the windows contain a constant number of SNPs, they will vary in physical length, as expressed in base pairs. This means that given two genomic windows $W_1$ and $W_2$, corresponding to different physical lengths, a different number of $S^*$ scores will be calculated. In order to ensure that all windows have the same number of features to feed into the model, we take the mean, standard deviation, and maximum of the $S^*$ score. Similarly, we supply the mean, standard deviation, and maximum of the non-outgroup SNP counts. These six values are then used as features for the model. These features are not included in the input matrix $G_W$, and instead are appended to intermediate embeddings before the final classification step of the model (Figure 1).

We use a haplotype-based adaptation of the $S^*$ statistic[6], calculating scores separately for each phased chromosome rather than at the diploid individual level. The statistic identifies chains of alleles absent from the outgroup population that are present in the target haplotype, with higher scores corresponding to a higher probability of the haplotype segment being of archaic origin.

### S6. iHS calculations

To identify signals consistent with positive selection, we relied on a haplotype-based test for selection by computing integrated haplotype scores (iHS)[7]. iHS measures the decay in linkage disequilibrium from a core SNP due to new mutations and recombination events, as recent positive selection is expected to result in long, frequent haplotypes with high haplotype homozygosity in a population. Normalized $|\text{iHS}| > 2$ reflects that the haplotype is longer and at a higher frequency than expected under neutrality, and is commonly considered the threshold for evidence of positive selection[8][7].

For each admixed population, we first computed the unstandardized iHS for each SNP with a minor allele frequency $> 0.05$ using `selscan v3.0.0`[8] and population specific recombination maps from Spence and Song (2019)[9] (https://zenodo.org/records/11437540), where ancestral states were polarized using a primate alignment as defined by the Enredo, Pecan, Ortheus (EPO) pipeline[10] (https://ftp.ensembl.org/pub/release-86/fasta/ancestral_alleles). These unstandardized values were then subsequently normalized in 100 evenly spaced derived allele frequency bins across the entire genome as described by Szpiech and Hernandez (2014)[8].

When reporting candidate genes for adaptive introgression in Figure 5, we sort genes vertically by taking the maximum normalized iHS observed for each gene across the four focal populations.
